## Supplementary material for "Comparative neuroimaging of the carnivoran brain: Neocortical sulcal anatomy"

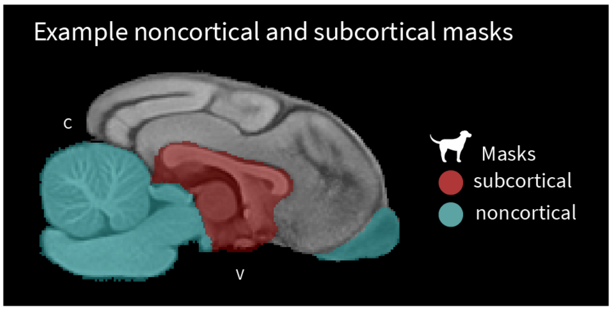


**Supplementary Figure S1. Illustration of manually created subcortical and noncortical masks for surface reconstruction.** For surface creation with precon_all, we created a subcortical (red) and noncortical (cyan) mask for each animal using ITK-snap (Yushkevich et al., 2006). The subcortical masks include the corpus callosum and expanded to the outer borders of the lateral ventricles; areas included in this mask are filled during surface generation. The noncortical masks include the olfactory bulb, brain stem and cerebellum to remove from the surface. The masks shown were created for the canine stereotactic breed-averaged template (Johnson et al., 2020). C, caudal; V, ventral.

### Supplementary Note 1: Major sulci from ventral perspective

The majority of the sulci seen from a ventral view (**Supplementary Figure S2**) were already described in the main text, as they were visible from lateral and dorsal views (see Figures **2-3**).

The lateral rhinal fissure represents the ventral border of the neocortex in carnivorans. The pseudo-sylvian fissure originates from the lateral rhinal fissure, dividing it into a caudal (light green; **Supplementary Figure S2**) and rostral (light blue; **Supplementary Figure S2**) part. The sulcal anatomy of the brown bear stood out compared to the other species as the caudal portion of the caudal lateral rhinal fissure merged with the retrosplenial sulcus (yellow; **Supplementary Figure**; and see **Supplementary Note 2**), and the marginal sulcus (**Supplementary Figure S2,** cyan) extended to the ventral surface terminating closely to the rostral end of the caudal lateral rhinal fissure. In all other species, the retrosplenial did not merge with the caudal lateral rhinal fissure, and the marginal sulcus was not visible on the ventral surface as it terminated more dorsally.

The ventral lateromedial sulcus (**Supplementary Figure S2,** pink) is only visible from a ventral perspective and is orthogonal to the occipitotemporal (red) and retrosplenial sulcus (yellow; see also **Supplementary Note 2**) positioned between the two sulci and the rostral lateral rhinal fissure (**Supplementary Figure S2**, light green). We were able to identify the ventral lateromedial sulcus in the Asian lion, brown and Asian-small clawed otter; and in most Canidae, except for the domestic dog and fennec fox. This sulcus has been previously described as not consistently identifiable in domestic dogs (Czeibert et al., 2018).

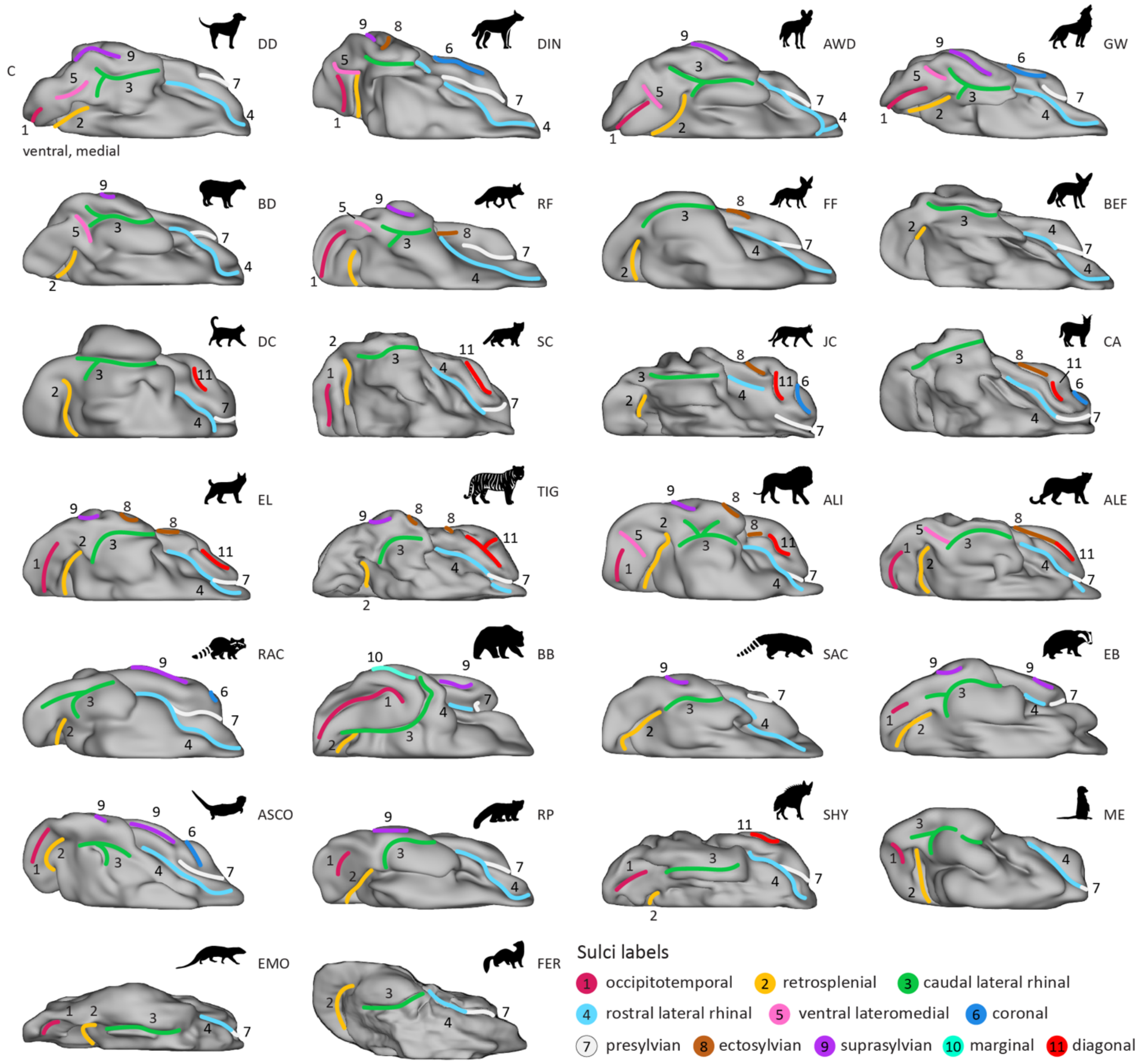


**Supplementary Figure S2. Ventral view of major neocortical sulci.** The rostral and caudal rhinal fissure represent the ventral border of the carnivoran neocortex. The majority of the sulci visible on the ventral view can also be seen from a dorsal and/or lateral perspective (**Figures 2-3**), and the retrosplenial and occipitotemporal sulcus extend ventrally from the medial wall (see **Supplementary Figure S4**). The ventral lateromedial sulcus (5, pink) can only be seen on the ventral surface of the occipital lobe. We were not able to consistently identify this sulcus in all Carnivoran species. Animal acronyms: ALI, Asiatic lion; ALE, Amur leopard; ASCO, Asian small-clawed otter; AWD, African wild dog; BB, brown bear; BD, bush dog; BEF, bat-eared fox; CA, caracal; DC, domestic cat; DD, domestic dog; DIN, dingo; EB, Eurasian badger; EL, Eurasian lynx; EMO, Egyptian mongoose; FER, ferret; FF, fennec fox; GW, grey wolf; JC, jungle cat; ME, meerkat; RAC, raccoon; RF, red fox; RP, red panda; SAC, South American coati; SC, sand cat; SHY, striped hyaena; TIG, Bengal tiger.

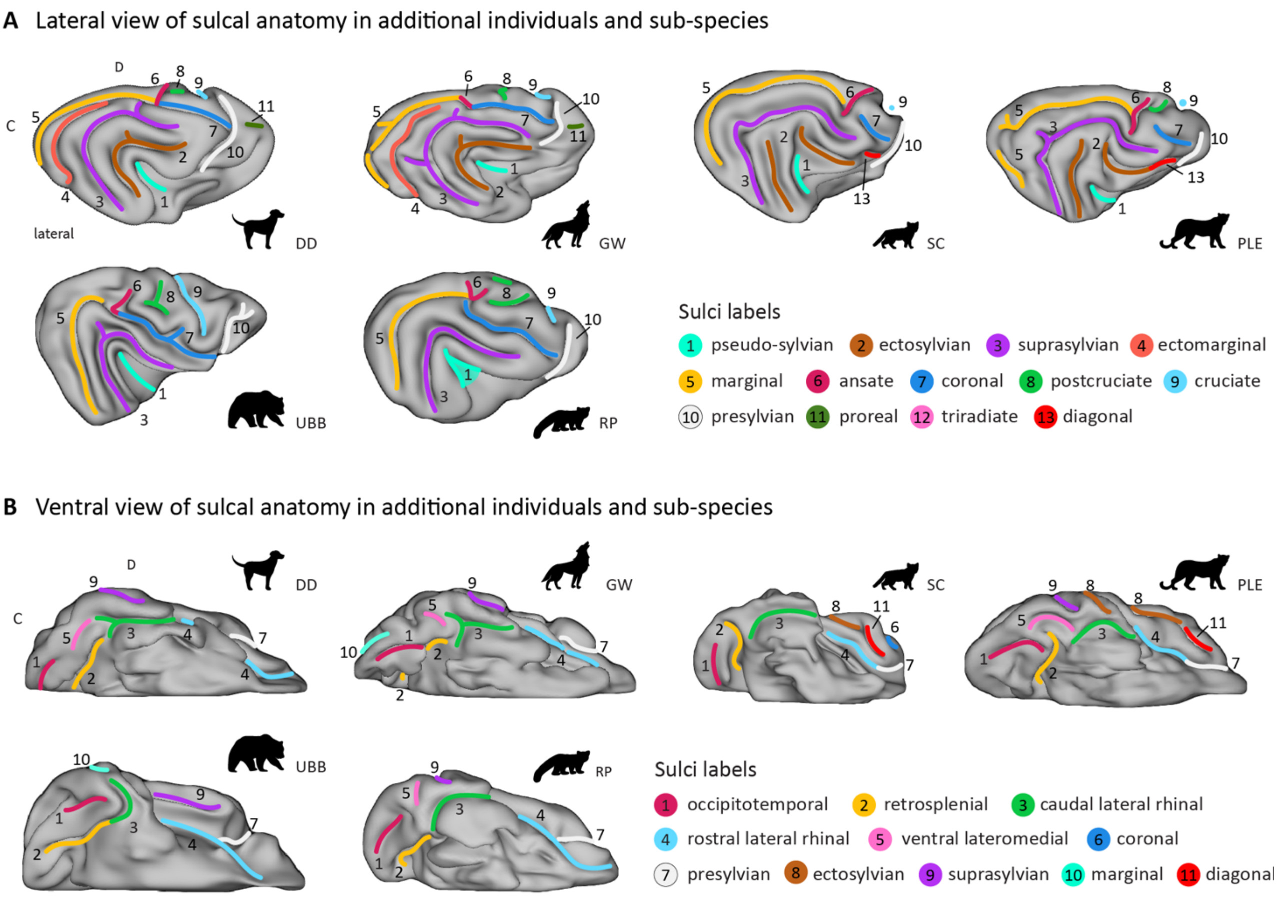


**Supplementary Figure S3. Lateral and medial view of major neocortical sulci in additional individuals and sub-species.** The sulcal topology of the additional individuals closely resembled the surfaces presented in the main text, with all major sulci present (see **Figure 2**, **Supplementary Figure S2**). Only minor variations in sulcal shape were observed. (**A**) For example, u-shaped pseudo-sylvian fissure (cyan, 1) of the second red panda was continuous in contrast to the split configuration observed in the other specimen (**Figure 2**). (**B**) Similar to both brown bear specimens, the marginal sulcus of the second grey wolf was also visible from the ventral view. In contrast, this sulcus was not visible in the ventral view of the first grey wolf or in any of the other species examined. Animal acronyms: DD, domestic dog; GW, grey wolf; SC, sand cat; PLE, Persian leopard; UBB, Ussuri brown bear; RP, red panda.

### Supplementary Note 2: Major sulci of the medial wall

The cruciate sulcus (**Supplementary Figure S4**, light blue) was also visible on the medial wall and merged in the majority of the species with the dorsal longitudinal splenial sulcus (**Supplementary Figure S4**, purple red). The splenial sulcus runs caudally and curves ventrally at the caudal end. The portion of the sulcus after the bend is called the retrosplenial sulcus (**Supplementary Figure S4, S5B**, yellow). In most of the brains in the present sample, the cruciate, splenial and retrosplenial sulci merged, or two of the three sulci merged; only in the brown bear and Ussuri brown bear all three sulci were detached.

In the wolf-like Canidae (the domestic dogs, dingo, African wild dog, and grey wolves), the Eurasian lynx, sand cats, Bengal tiger, Asiatic lion, striped hyaena and both the Amur and Persian leopard, we noted another sulcus curving around the caudal bend of the retro-

/splenial sulcus. The dorsal portion of the sulcus is called the suprasplenial sulcus (**Supplementary Figure S4, S5B**, green), and the ventral portion is the occipitotemporal sulcus (**Supplementary Figure S2-5B**, purple). In the dingo, Bengal tiger, Asiatic lion, striped hyena and the Amur and Persian leopard, the suprasplenial and the occipitotemporal sulcus were detached. In most other species, only one of the two sulci could be identified, and in the domestic cat, fennec fox, South American coati, and ferret, neither was observed.

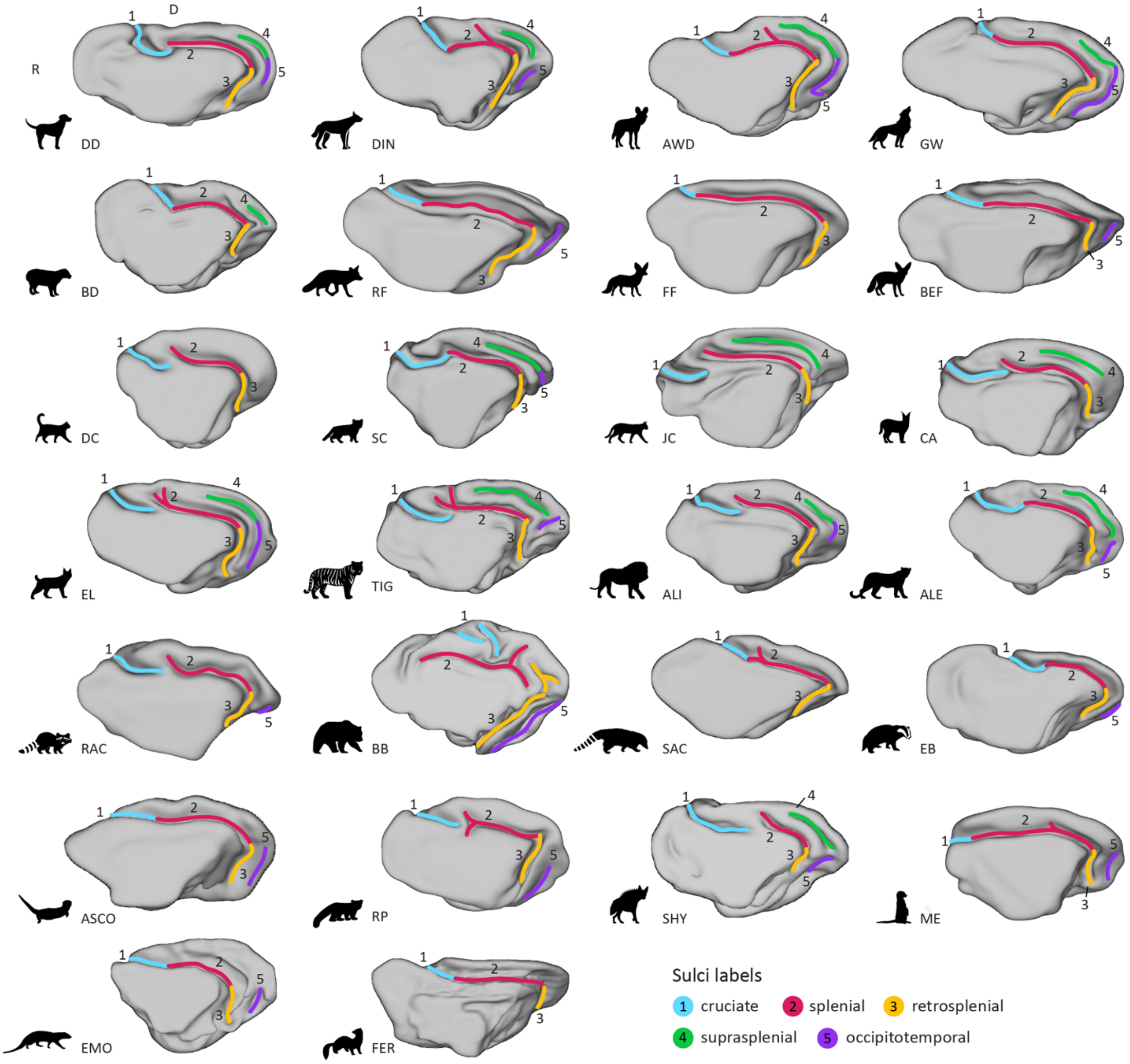


**Supplementary Figure S4. Major neocortical sulci in the medial wall.** The cruciate sulcus (light blue, 1) was the only sulcus that was also visible on the lateral and dorsal view (see **Figures 2-3**). All species had a cruciate, splenial (purple red, 2), and retrosplenial (yellow, 3) sulcus. Occurrence and appearance of the suprasplenial (green, 4) and occipitotemporal (purple, 5) sulci varied across species. Anatomical locations are indicated on the domestic dog surface in the top row, left corner. R, rostral; D, dorsal. Animal acronyms: ALI, Asiatic lion; ALE, Amur leopard; ASCO, Asian small-clawed otter; AWD, African wild dog; BB, brown bear; BD, bush dog; BEF, bat-eared fox; CA, caracal; DC, domestic cat; DD, domestic dog; DIN, dingo; EB, Eurasian badger; EL, Eurasian lynx; EMO, Egyptian mongoose; FER, ferret; FF, fennec fox; GW, grey wolf; JC, jungle cat; ME, meerkat; RAC, raccoon; RF, red fox; RP, red panda; SAC, South American coati; SC, sand cat; SHY, striped hyaena; TIG, Bengal tiger.

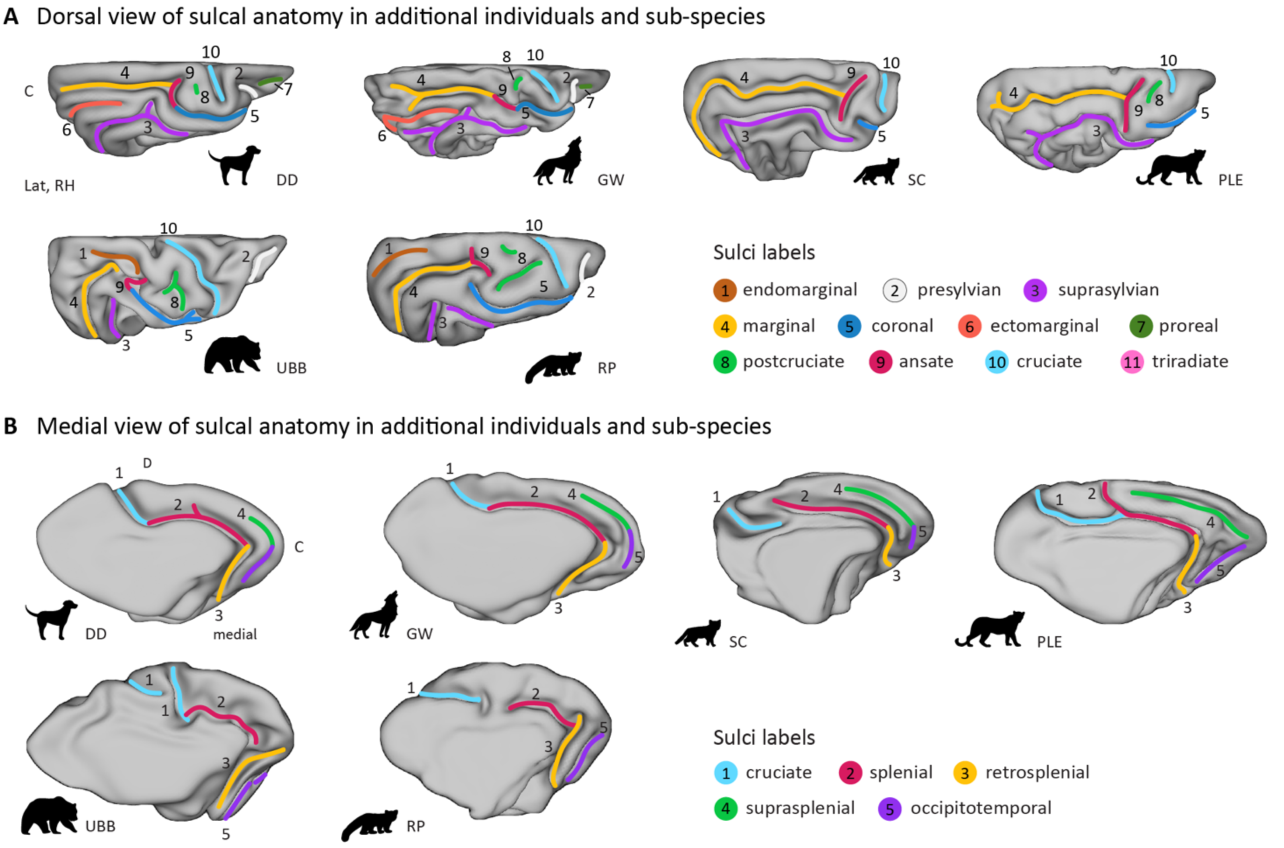


**Supplementary Figure S5. Dorsal and medial view of major neocortical sulci in additional individuals and sub-species.** None of the supplementary samples revealed major variations in shape or presence of the sulcus; only minor variations in sulcal shape were observed. (**A**) For example, in the Ussuri brown bear (UBB), the ansate sulcus (purple, 9) was more clearly identifiable than in the brown bear, as it displayed the typical perpendicular orientation relative to the marginal sulcus. (**B**) In the second sand cat, the cruciate (light blue, 1) and splenial (purple red, 2) sulci were detached, a variation also observed in several other species (**Supplementary Figure S4**) whereas in the first sand cat, these sulci were merged. Animal acronyms: DD, domestic dog; GW, grey wolf; SC, sand cat; PLE, Persian leopard; UBB, Ussuri brown bear; RP, red panda.

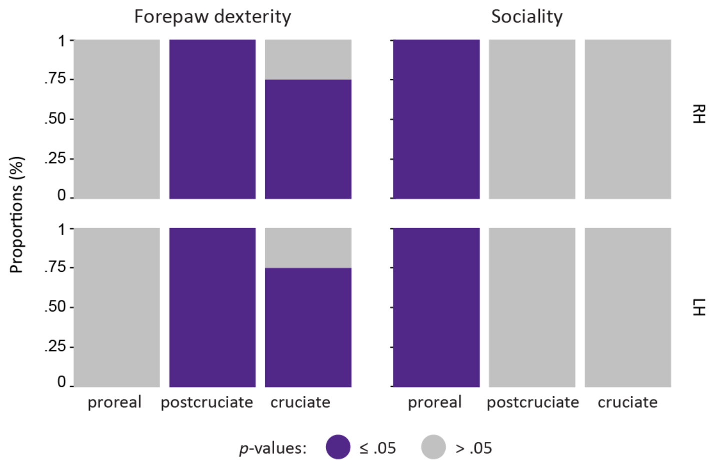


**Supplementary Figure S6. Proportion of significant associations between relative sulcal length and behavioural characteristics.** Preliminary quantitative analyses, conducted to complement qualitative descriptions (**Figure 5**), reveal that the relative length of the proreal sulcus was consistently greater in cooperatively hunting species than in solitary hunters regardless of choice for reference sulcus ($n$ = 7) and both hemispheres, while no effect of forepaw dexterity was observed (**Supplementary Table S1**). In contrast, analyses of the postcruciate sulcus revealed the opposite pattern, with consistently greater sulcal length in species with high forepaw dexterity across all reference sulci ($n$ = 4) and in both hemispheres, and no signification relationship with sociality, (**Supplementary Table S2**). Analyses of the cruciate sulcus length ($n$ = 4 reference sulci) revealed a positive relationship with high forepaw dexterity in 75% of models in each hemisphere, but no significant relationship with sociality (**Supplementary Table S3**). RH, right hemisphere; LH, left hemisphere.

**Supplementary Table S1**. Effects of forepaw dexterity and sociality on relative length of the proreal sulcus

| Predictor | Reference sulcus | Hemisphere | $F$ ($df_{num}$, $df_{den}$) | $p$ | $\eta^2_p$ |
|---|---|---|---|---|---|
| Forepaw dexterity | ansate | right | 1.23 (1, 22) | .28030 | .13 |
| Sociality | ansate | right | 25.43 (1, 22) | **< .00001** | .54 |
| Forepaw dexterity | coronal | right | 1.57 (1, 22) | .22380 | .16 |
| Sociality | coronal | right | 32.53 (1, 22) | **< .00001** | .60 |
| Forepaw dexterity | marginal | right | 1.39 (1, 22) | .25050 | .15 |
| Sociality | marginal | right | 28.91 (1, 22) | **< .00001** | .57 |
| Forepaw dexterity | presylvian | right | 1.75 (1, 22) | .19890 | .18 |
| Sociality | presylvian | right | 36.42 (1, 22) | **< .00001** | .62 |
| Forepaw dexterity | retrosplenial | right | 1.61 (1, 22) | .21740 | .17 |
| Sociality | retrosplenial | right | 33.47 (1, 22) | **< .00001** | .60 |
| Forepaw dexterity | splenial | right | 1.72 (1, 22) | .20370 | .18 |
| Sociality | splenial | right | 35.62 (1, 22) | **< .00001** | .62 |
| Forepaw dexterity | suprasylvian | right | 1.53 (1, 22) | .22880 | .16 |
| Sociality | suprasylvian | right | 31.81 (1, 22) | **< .00001** | .59 |
| Forepaw dexterity | ansate | left | .78 (1, 22) | .38580 | .09 |
| Sociality | ansate | left | 16.25 (1, 22) | **.00060** | .42 |
| Forepaw dexterity | coronal | left | .87 (1, 22) | .36220 | .10 |
| Sociality | coronal | left | 17.97 (1, 22) | **.00030** | .45 |
| Forepaw dexterity | marginal | left | .88 (1, 22) | .35900 | .10 |
| Sociality | marginal | left | 18.22 (1, 22) | **.00030** | .45 |
| Forepaw dexterity | presylvian | left | .92 (1, 22) | .34820 | .10 |
| Sociality | presylvian | left | 19.07 (1, 22) | **.00020** | .46 |
| Forepaw dexterity | retrosplenial | left | .58 (1, 22) | .45290 | .07 |
| Sociality | retrosplenial | left | 12.12 (1, 22) | **.00210** | .36 |
| Forepaw dexterity | splenial | left | .93 (1, 22) | .34610 | .10 |
| Sociality | splenial | left | 19.24 (1, 22) | **.00020** | .47 |
| Forepaw dexterity | suprasylvian | left | .91 (1, 22) | .35070 | .10 |
| Sociality | suprasylvian | left | 18.87 (1, 22) | **.00030** | .46 |

*Note*. Linear models were used to test the effects of two categorical behavioural predictors: forepaw dexterity (low, high) and sociality (solitary, cooperative hunting) on the relative length of the proreal sulcus. Models were fit separately for each hemisphere (left, right) and each target-reference sulcus pair. Significant *p*-values (bolded) indicate greater relative sulcal length in species with high dexterity or cooperative hunting behaviour. Proportions of significant results are presented in **Supplementary Figure S6**. P $df_{num}$, degrees of freedom numerator; $df_{den}$ degrees of freedom denominator; $\eta^2_p$, partial eta squared

**Supplementary Table S2**. Effects of forepaw dexterity and sociality on relative length of the postcruciate sulcus

| Predictor | Reference sulcus | Hemisphere | $F$ ($df_{num}$, $df_{den}$) | $p$ | $\eta^2_p$ |
|---|---|---|---|---|---|
| Forepaw dexterity | marginal | right | 71.71 (1, 22) | **< .00001** | .75 |
| Sociality | marginal | right | 5.28 (1, 22) | **.03150** | .19 |
| Forepaw dexterity | retrosplenial | right | 46.79 (1, 22) | **< .00001** | .66 |
| Sociality | retrosplenial | right | 3.94 (1, 22) | .05970 | .15 |
| Forepaw dexterity | splenial | right | 56.22 (1, 22) | **< .00001** | .70 |
| Sociality | splenial | right | 4.09 (1, 22) | .05560 | .16 |
| Forepaw dexterity | suprasylvian | right | 52.06 (1, 22) | **< .00001** | .69 |
| Sociality | suprasylvian | right | 4.72 (1, 22) | **.04080** | .18 |
| Forepaw dexterity | marginal | left | 38.24 (1, 22) | **< .00001** | .64 |
| Sociality | marginal | left | .00 (1, 22) | .97190 | .00 |
| Forepaw dexterity | retrosplenial | left | 30.57 (1, 22) | **< .00001** | .59 |
| Sociality | retrosplenial | left | .00 (1, 22) | .96130 | .00 |
| Forepaw dexterity | splenial | left | 84.59 (1, 22) | **< .00001** | .79 |
| Sociality | splenial | left | .61 (1, 22) | .44400 | .03 |
| Forepaw dexterity | suprasylvian | left | 96.33 (1, 22) | **< .00001** | .82 |
| Sociality | suprasylvian | left | .04 (1, 22) | .84340 | .00 |

*Note*. Linear models were used to test the effects of two categorical behavioural predictors: forepaw dexterity (low, high) and sociality (solitary, cooperative hunting) on the relative length of the postcruciate sulcus. Models were fit separately for each hemisphere (left, right) and each target-reference sulcus pair. Significant *p*-values (bolded) indicate greater relative sulcal length in species with high dexterity or cooperative hunting behaviour. Proportions of significant results are presented in **Supplementary Figure S6**. P $df_{\mathrm{num}}$, degrees of freedom numerator; $df_{\mathrm{den}}$ degrees of freedom denominator; $\eta^2_p$, partial eta squared

**Supplementary Table S3**. Effects of forepaw dexterity and sociality on relative length of the cruciate sulcus

| Predictor | Reference sulcus | Hemisphere | $F$ ($df_{num}$, $df_{den}$) | $p$ | $\eta^2_p$ |
|---|---|---|---|---|---|
| Forepaw dexterity | marginal | right | 7.24 (1, 22) | **.01330** | .25 |
| Sociality | marginal | right | .00 (1, 22) | .96900 | .00 |
| Forepaw dexterity | retrosplenial | right | 1.45 (1, 22) | .24210 | .09 |
| Sociality | retrosplenial | right | 2.36 (1, 22) | .13870 | .10 |
| Forepaw dexterity | splenial | right | 5.21 (1, 22) | **.03240** | .22 |
| Sociality | splenial | right | 1.31 (1, 22) | .26500 | .06 |
| Forepaw dexterity | suprasylvian | right | 7.63 (1, 22) | **.01130** | .28 |
| Sociality | suprasylvian | right | 1.09 (1, 22) | .30680 | .05 |
| Forepaw dexterity | marginal | left | 12.15 (1, 22) | **.00210** | .37 |
| Sociality | marginal | left | .12 (1, 22) | .72980 | .01 |
| Forepaw dexterity | retrosplenial | left | .80 (1, 22) | .38200 | .05 |
| Sociality | retrosplenial | left | 2.11 (1, 22) | .16080 | .09 |
| Forepaw dexterity | splenial | left | 8.19 (1, 22) | **.00910** | .29 |
| Sociality | splenial | left | .32 (1, 22) | .57990 | .01 |
| Forepaw dexterity | suprasylvian | left | 19.41 (1, 22) | **.00020** | .50 |
| Sociality | suprasylvian | left | 3.06 (1, 22) | .09390 | .12 |

*Note*. Linear models were used to test the effects of two categorical behavioural predictors: forepaw dexterity (low, high) and sociality (solitary, cooperative hunting) on the relative length of the cruciate sulcus. Models were fit separately for each hemisphere (left, right) and each target-reference sulcus pair. Significant *p*-values (bolded) indicate greater relative sulcal length in species with high dexterity or cooperative hunting behaviour. Proportions of significant results are presented in **Supplementary Figure S6**. P $df_{num}$, degrees of freedom numerator; $df_{den}$ degrees of freedom denominator; $\eta^2_p$, partial eta squared.
